# ASCL1-ERK1/2 Axis: ASCL1 restrains ERK1/2 via the dual specificity phosphatase DUSP6 to promote survival of a subset of neuroendocrine lung cancers

**DOI:** 10.1101/2023.06.15.545148

**Authors:** Ana Martin-Vega, Svetlana Earnest, Alexander Augustyn, Chonlarat Wichaidit, Adi Gazdar, Luc Girard, Michael Peyton, Rahul K. Kollipara, John D. Minna, Jane E. Johnson, Melanie H. Cobb

**Author notes:** Equal contribution (Ana Martin-Vega, Svetlana Earnest, Alexander Augustyn). University of Texas MD Anderson Cancer Center, Department of Radiation Oncology, Houston, TX.

## Abstract

The transcription factor achaete-scute complex homolog 1 (ASCL1) is a lineage oncogene that is central for the growth and survival of small cell lung cancers (SCLC) and neuroendocrine non-small cell lung cancers (NSCLC-NE) that express it. Targeting ASCL1, or its downstream pathways, remains a challenge. However, a potential clue to overcoming this challenage has been information that SCLC and NSCLC-NE that express ASCL1 exhibit extremely low ERK1/2 activity, and efforts to increase ERK1/2 activity lead to inhibition of SCLC growth and surival. Of course, this is in dramatic contrast to the majority of NSCLCs where high activity of the ERK pathway plays a major role in cancer pathogenesis. A major knowledge gap is defining the mechanism(s) underlying the low ERK1/2 activity in SCLC, determining if ERK1/2 activity and ASCL1 function are inter-related, and if manipulating ERK1/2 activity provides a new therapeutic strategy for SCLC. We first found that expression of ERK signaling and ASCL1 have an inverse relationship in NE lung cancers: knocking down ASCL1 in SCLCs and NE-NSCLCs increased active ERK1/2, while inhibition of residual SCLC/NSCLC-NE ERK1/2 activity with a MEK inhibitor increased ASCL1 expression. To determine the effects of ERK activity on expression of other genes, we obtained RNA-seq from ASCL1-expressing lung tumor cells treated with an ERK pathway MEK inhibitor and identified down-regulated genes (such as SPRY4, ETV5, DUSP6, SPRED1) that potentially could influence SCLC/NSCLC-NE tumor cell survival. This led us to discover that genes regulated by MEK inhibition suppress ERK activation and CHIP-seq demonstrated these are bound by ASCL1. In addition, SPRY4, DUSP6, SPRED1 are known suppressors of the ERK1/2 pathway, while ETV5 regulates DUSP6. Survival of NE lung tumors was inhibited by activation of ERK1/2 and a subset of ASCL1-high NE lung tumors expressed DUSP6. Because the dual specificity phosphatase 6 (DUSP6) is an ERK1/2-selective phosphatase that inactivates these kinases and has a pharmacologic inhibitor, we focused mechanistic studies on DUSP6. These studies showed: Inhibition of DUSP6 increased active ERK1/2, which accumulated in the nucleus; pharmacologic and genetic inhibition of DUSP6 affected proliferation and survival of ASCL1-high NE lung cancers; and that knockout of DUSP6 “cured” some SCLCs while in others resistance rapidly developed indicating a bypass mechanism was activated. Thus, our findings fill this knowledge gap and indicate that combined expression of ASCL1, DUSP6 and low phospho-ERK1/2 identify some neuroendocrine lung cancers for which DUSP6 may be a therapeutic target.

## Introduction

Increased expression of the basic helix-loop-helix transcription factor achaete-scute complex homolog 1 (ASCL1; also referred to as HASH1, MASH1) was observed in neuroendocrine lung tumors more than 30 years ago^1^. ASCL1 expression is also found in prostate tumors, glioblastoma, neuroblastoma, medullary thyroid cancer, thymic carcinoma^2–5^ and other cancers with neuroendocrine characteristics including the majority of small cell lung cancers (SCLC) and nonsmall cell lung cancers with neuroendocrine features (NSCLC-NE)^6, 7, 8–15, 16^. Depletion of ASCL1 from patient-derived SCLC lines reduced the formation of colonies in soft agar and reduced growth of xenograft tumors in nude mice^6, 7, 15, 16^. Conditional disruption of *Ascl1* in the lung prevented formation of tumors in a genetically engineered mouse model (GEMM) of neuroendocrine lung cancer^17^. These findings support the conclusion that ASCL1 is an oncogenic driver on which certain neuroendocrine lung cancers depend for survival and continued growth.

ASCL1 is a lineage transcription factor required for neuronal differentiation^18, 19^ and for an early stage of differentiation in specific neuroendocrine lineages including neuroendocrine cells in the lung and adrenal medulla^20, 21^. ASCL1 is essential during the development of the lineages in which it is expressed. In adult tissues, it is restricted to stem or progenitor cell niches such as in the brain^22–24^. Accordingly, in the adult lung, it is restricted to the rare neuroendocrine bodies^25–27^. Deletion of mouse *Ascl1* from the pulmonary neuroendocrine lineage demonstrated that ASCL1 is required for normal development of this lung cell type^27–31^.

In general, an unusually high number of genetic alterations have been identified in SCLCs with common loss of tumor suppressor genes, TP53 and RB1, high expression of myc family members, and increased copy number and mutations in other transcription factors^14, 32–34^. In many NSCLCs, the activities of the mitogen-activated protein kinases (MAPKs) ERK1/2 are elevated due to the occurrence of mutations in upstream molecules (oncogenes) that activate this pathway. Mutations are common in tyrosine kinases such as the EGF receptor (EGFR) and fusion mutations of ALK^35^. Mutations in *KRAS* and *BRAF* genes are also common in NSCLC including those with neuroendocrine characteristics. In contrast, in SCLC, which account for approximately 15% of newly diagnosed lung cancers^36, 37^, mutations in these oncogenes are rare to nonexistent. Recent work by Rubio *et al.* also highlighted the importance of ERK activation in triggering some SCLC subtypes, as they observed in one NeuroD1-high (NCI-H82) and one YAP1-high (NCI-H196) tumor cell line. Of interest, they found that the integrin subunit ITGB2 was crucial for EGFR activation and tumorigenesis in these two cell lines^38^.

Twenty-five years ago, expression of a Raf-1 truncation fused to the hormone binding domain of the estrogen receptor was used to artificially induce ERK1/2 activation in an estradiol-dependent manner to test the impact of activation of ERK1/2 on two SCLC lines ^39, 40^. Elevated ERK1/2 activity caused growth arrest with an accumulation of cells in G1, induction of p27 and p16, proteins that cause cell cycle arrest, and reduced expression of neuroendocrine markers including secreted peptides. These studies suggested that the activities of ERK1/2 interfere with SCLC growth, in spite of the fact that ERK1/2 are thought to have essential functions in disease progression. In fact, two recent studies have shown a signficant negative correlation between ERK activity and ASCL1 expression^41, 42^. Thus, questions have been raised about the importance of ERK1/2 to tumorigenicity of neuroendocrine lung cancers and the therapeutic potential of this pathway^43–47^. Despite the aforementioned ERK negative impact in SCLC growth, it is important to note that these cells still rely on ERK activation, albeit lower than observed in NSCLC, to survive. In a parallel project, we have not been able to simultaneously silence both ERK1 and ERK2 isoforms (unpublished data), which indicates a low but essential optimal ERK activity.

To explore the relevance of these findings in multiple neuroendocrine lung cancers, we examined the interplay between ASCL1 and ERK1/2 activities and the consequences of stimulating and inhibiting the ERK pathway on cell survival. We noted that the dual specificity MAP kinase phosphatase, DUSP6 (also known as MKP3), is expressed in subsets of SCLC and NSCLC-NE lines expressing high levels of ASCL1 (“ASCL1-high”). DUSP6 is a phosphatase with selectivity for ERK1/2 over other MAPK family members^48–51^. Because ERK1/2 activity negatively impacts SCLC growth and survival, we tested the possibility that function of DUSP6 (by dephosphorylating and inactivating ERK1/2) was necessary for the survival of these SCLC and NSCLC-NE lines and thus, a potential therapeutic target.

## Results

### Expression of ERK signaling and ASCL1 have an inverse relationship in NE lung cancer models

Because earlier studies suggested that ERK activity is harmful to neuroendocrine lung cancer cells^39, 40^, we asked whether the function of ASCL1 is responsible for suppressing the ERK pathway. ASCL1 was knocked down and activation of ERK1/2 was assessed in the NSCLC-NE, ASCL1 high HCC1833 cells (Figure 1A). We found that reduced ASCL1 expression ih HCC1833 cells resulted in increased ERK1/2 activity, which was detected using antibodies that recognize their dually phosphorylated, active forms (pERK2, pT183, pY185). We then examined the converse, and tested the consequences of suppressing ERK activity on ASCL1 expression. ERK activation was prevented using an inhibitor, PD0325901 (PD03), of the upstream protein kinases MEK1/2. Exposure of HCC1833 cells to PD03 caused an increase in the accumulation of ASCL1 protein, indicating an inverse relationship between ERK activity and ASCL1 protein levels. Notably, the half-life of ASCL1 protein in HCC1833 NSCLC-NE is short (<15 minutes) as measured by treating cells with cycloheximide to prevent new protein synthesis, and there was no difference in its rate of decay when ERK activity was decreased using the MEK inhibitor PD03 (Figure 1B, Figure S1A-C). To determine if this effect was due to increased ASCL1 mRNA, we analyzed *ASCL1* mRNA and found it increased in cells in which ERK1/2 activation was blocked with the MEK inhibitor PD03. This increase in *ASCL1* mRNA was prevented by the inclusion of actinomycin D to inhibit new mRNA synthesis (Figure 1C). These studies indicate that ERK1/2 have little or no effect on the stability of ASCL1 protein but decrease *ASCL1* gene expression. Together, these results suggest that ERK1/2 signaling interferes with ASCL1 expression and vice versa, identifying an ERK/ASCL1 axis.

**Figure 1.**
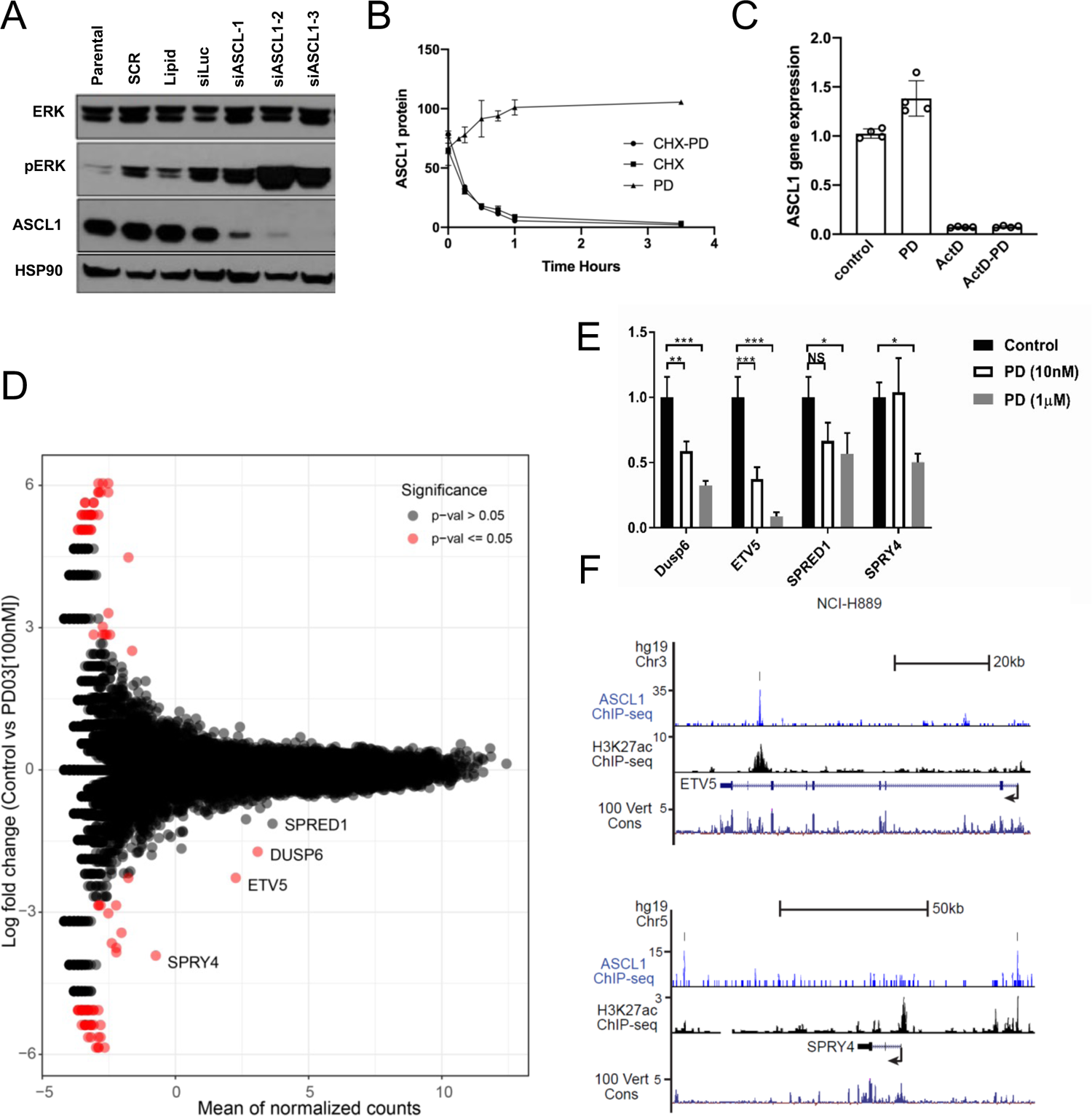
Interplay of ERK1/2 and ASCL1 in neuroendocrine lung cancer cells. **A.** ASCL1 was depleted from HCC1833 cells using three different siRNAs and compared to a series of controls omitting siASCL1 oligonucleotides. Total ERK (ERK1, upper band; ERK2 lower band) and phosphorylated ERK (pERK, pT185, pY185) were immunoblotted, as was ASCL1. HSP90 was immunoblotted as a loading control. **B.** HCC1833 cells were treated with 1 μM PD0325901 or DMSO control prior to the addition of 128 μM cycloheximide and cells were harvested over a 3.5 hour time course. Two biological replicates. ASCL1 protein quantified by LiCOR imaging of immunoblots with guinea pig anti-ASCL1 antibodies (see Figure S1A for immunoblots) is plotted over time. **C.** HCC1833 cells were treated with 1 µM PD0325901 or DMSO control prior to the addition of actinomycin D. ASCL1 mRNA was measured by quantitative RT-PCR in two replicate time courses. ASCL1 mRNA comparing PD03 in the absence of actinomycin D, P = 0.0009. **D.** RNA-seq was performed on H889 cells treated with 100 nM PD03 or DMSO control for 18 hours. Similar results were obtained with 1 μM PD03 (see Table S1). Red dots indicate genes most significantly altered by PD03. **E.** Effects of PD03 in H889 cells on expression of 4 genes measured by quantitative RT-PCR. N=3. **F.** Occupancy of the ETV5 gene (top) and the SPRY4 gene (bottom) by ASCL1 extracted from ChIP-seq data from H889 cells. H3K27Ac for each of these gene regions is also shown. 100 Vert Cons, the last line in each set, is the sequence conservation across 100 vertebrates.

### Genes regulated by MEK inhibition suppress ERK activation and CHIP-seq reveals that these are bound by ASCL1

To gain mechanistic insight into the interplay between ERK signaling and ASCL1, we performed RNA-seq in a SCLC cell line H889 in the absence and presence of PD03. Numerous changes in gene expression were detected but among those most decreased by blocking ERK pathway signaling were *DUSP6, SPRY4* and to a lesser extent *SPRED1* (Figure 1D); all three are known to suppress signaling by the ERK pathway. The dual specificity phosphatase DUSP6 is an ERK-selective phosphatase that directly inactivates ERK1/2 activity, while sprouty proteins such as SPRY4 and the related protein SPRED1 work upstream in the ERK pathway to prevent activation of the ERK kinase cascade at the receptor level^52–54^. Blocking ERK pathway activity apparently obviates the need to induce negative feedback via expression of these genes. Also suppressed by the MEK inhibitor was expression of the ETS family transcription factor *ETV5* (Figure 1D, Table S1). Interestingly ETV5 and other ETS factors have been linked to transcription of DUSP6^55–59^. Changes in expression of *DUSP6, SPRY4, ETV5,* and *SPRED1* were confirmed by quantitative RT-PCR in the H889 cells with and without PD03 (Figure 1E). Furthermore, we learned from chromatin immunoprecipitation-sequencing that ASCL1 is found on SPRY4 and ETV5 genes in H889 cells (Figure 1F), as well as, on DUSP6 in several ASCL1-high SCLC and NE-NSCLC cell lines (Figure 3B). We also evaluated the effects of a less potent MEK inhibitor (AZD6244) on survival of H889 and HCC1833 cell lines. We found little effect on survival of the SCLC cell line H889 (Figure S1D), as concentrations several orders of magnitude above the IC_50_ did not reduce survival, and a relatively smaller degree of insensitivity of HCC1833 (Figure S1E). These results are consistent with the reduced requirement of these cells for ERK1/2 activity.

### Survival of neuroendocrine lung tumor cells is inhibited by activation of ERK1/2

Previous research in a neuroblastoma cell line demonstrated that treating cells with phorbol 12-myristate 13-acetate (PMA), which acutely stimulates protein kinase C leading to activation of the ERK pathway, resulted in rapid degradation of *ASCL1* mRNA^60^. Therefore, we tested PMA as an alternative means to activate ERK1/2 and to determine its effects on *ASCL1* expression. Treating HCC1833 cells for 3 h with 100 nM PMA resulted in a time-dependent decrease of *ASCL1* mRNA (Figure 2A). Over the same time course, ERK1/2 were strongly activated (Figure 2B), particularly compared to serum stimulation of H889 (Figure S2A). Although PMA is an activator of ERK signaling, and increases ERK phosphorylation during short-term treatments, at high concentrations it also promotes PKC degradation. Predictably, following 24 h of PMA exposure, we detected very little pERK1/2 consistent with loss of the stimulatory PKCs by high concentrations of phorbol ester. Cell cycle analysis demonstrated a 10-fold induction of cell death at 24 h, even at 1 nM PMA, in contrast to effects on immortalized human bronchial epithelial cells, HBEC-3KT^61^, which are less sensitive to PMA (Figure 2C, D). After 24 h, all concentrations of PMA caused a loss of ASCL1 protein (Figure 2E). Additionally, an increase in cleaved PARP was identified, further suggesting that cell death occurs via apoptosis (Figure 2E). These results further support the suggestion of a negative relationship between ERK activation and ASCL1 expression.

**Figure 2.**
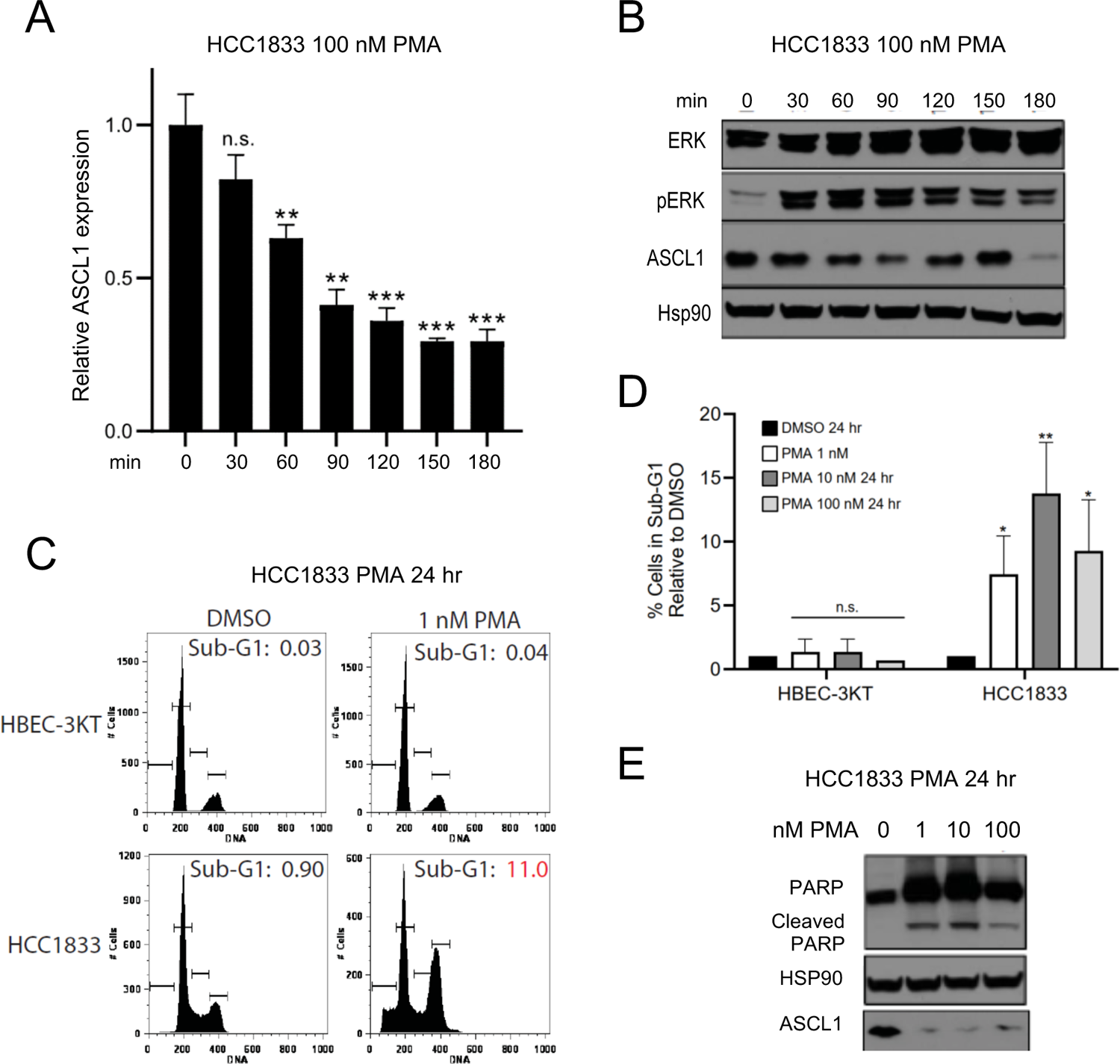
PMA activates ERK, suppresses ASCL1, and induces PARP cleavage. **A.** Short-term treatments with PMA were performed in 6-well plates on 4×10^5^ HCC1833 cells treated with 100 nM PMA for 30, 60, 90, 120, 150, and 180 min. Expression of ASCL1 was quantified by quantitative RT-PCR. Bar graphs represent the average of 3 similar experiments. P values **p<0.01, ***p<0.005. **B.** Over the same time course, ERK, pERK, and ASCL1 were immunoblotted, with HSP90 as the loading control. **C.** and **D.** Cell cycle analysis by flow cytometry of HCC1833 cells and immortalized normal-like HBEC-3KT cells treated with 1 nM PMA for 24 hours. **E.** HCC1833 cells were treated with 1, 10 or 100 nM PMA for 24 hours. Cleaved PARP and ASCL1 were detected by immunoblotting.

### A subset of ASCL1-high NSCLC-NE and SCLC cell lines express DUSP6

Because DUSP6 is a known regulator of ERK activity, we evaluated a larger group of neuroendocrine lung cancers for DUSP6 expression. RNA-seq data indicated much higher FPKM for *DUSP6* mRNA in ASCL1-expressing compared to SCLCs which highly express NEUROD1, another lineage transcription factor overexpressed and driving a subset of SCLCs^62^ (Figure 3A). As mentioned above, chromatin immunoprecipitation-sequencing of ASCL1-high (SCLC, red; NSCLC-NE, purple) and NEUROD1-high (blue) NSCLC-NE and SCLC lines^14, 17^ demonstrated that ASCL1 but not NEUROD1 was bound to an area just downstream of the *DUSP6* gene in a region with the active chromatin mark H3K27-acetylated (Figure 3B) in a number of SCLC and NSCLC-NE lines. The enrichment of acetylated H3K27 encompassing these ASCL1-bound regions, and the expression of DUSP6 in the SCLC-ASCL1 high and NE-NSCLC cells, suggest *DUSP6* is not only an ERK-regulated but also an ASCL1-regulated gene^17^.

**Figure 3.**
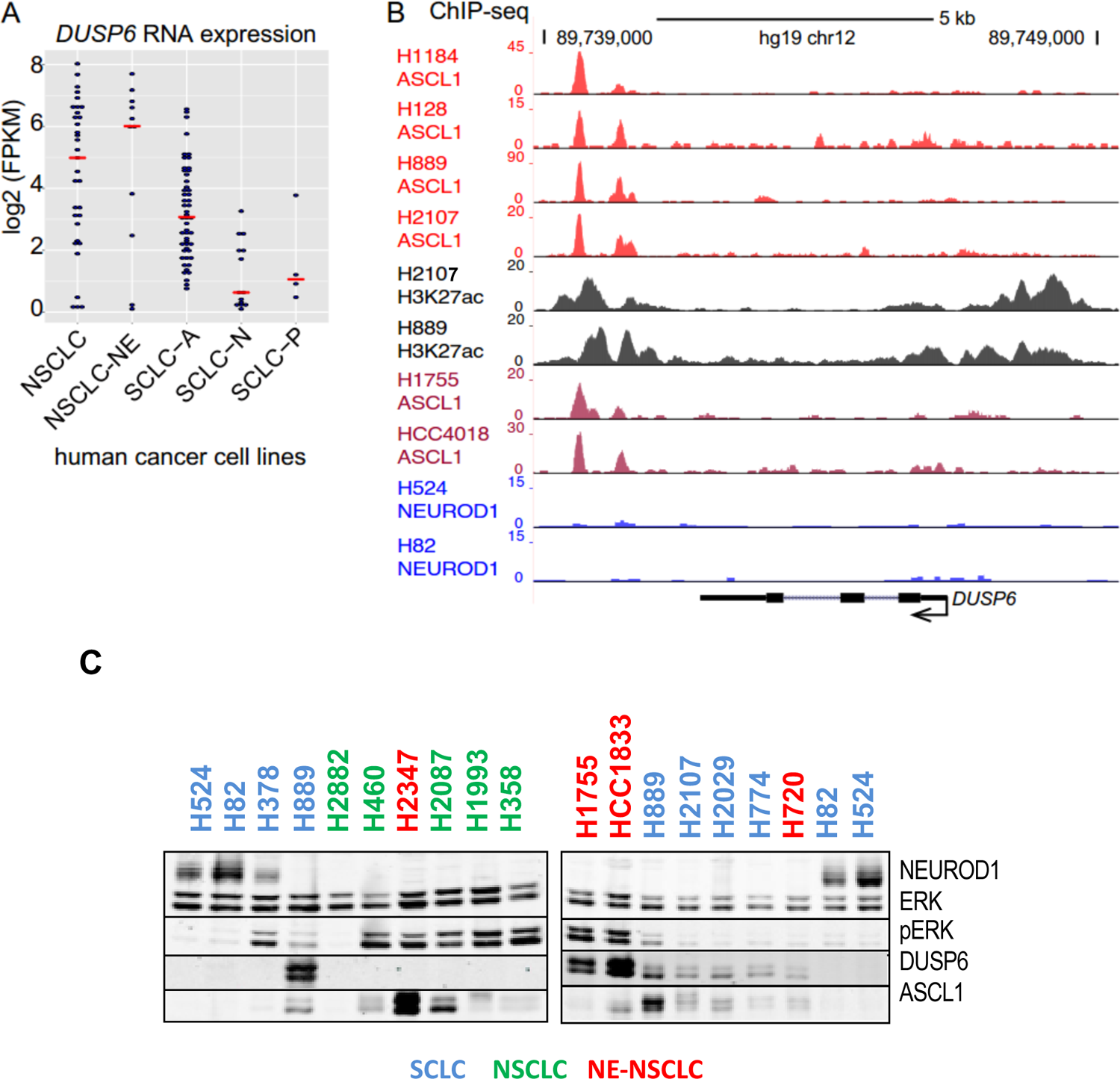
DUSP6 enrichment in ASCL1-expressing SCLC and NSCLC-NE and the DUSP6 gene is bound by ASCL1. **A.** The expression of mRNA encoding DUSP6 is shown for a large panel of lung cancer cell lines including NSCLC, NSCLC-NE, and three subsets of SCLC: ASCL1-high cells, SCLC-A; NEUROD1-high cells, SCLC-N; SCLC not expressing ASCL1 or NEUROD1, SCLC-P. **B.** ChIP-seq data from^17^ showing ASCL1, NEUROD1, and H3K27ac occupancy in the DUSP6 gene locus. Red, ASCL1 ChiP-seq in SCLC-A cell lines; magenta, ASCL1 ChiP-seq in NSCLC-NE cell lines; blue, NEUROD1 ChiP-seq in SCLC-N cell lines; black, H3K27ac ChIP-seq in SCLC-A cell lines. **C.** Immunoblotting of the indicated cell lines for ERK and pERK, ASCL1, NEUROD1, and DUSP6 is shown. SCLC (blue) - H524, H82, H889, H2107, H2029, H774; NE-NSCLC (red) - HCC2374, H1755, HCC1833; NSCLC (green) - H2882, H460, H2087, H1993, H358; carcinoid - H720. Total ERK is used as a loading control. See Table S2 for more complete descriptions.

To assess all of these key protein factors at one time we performed immunoblots for expression of ASCL1, NEUROD1, ERK, pERK, and DUSP6 (Figure 3C). Among lung cancer cell lines, ASCL1 was detected primarily in SCLC and NSCLC-NE. Immunoblotting of lysates from lung cancer lines, including some NSCLC-NE that are ASCL1-high (e.g. HCC1833 and HCC2374), showed that ERK1/2 were generally more highly phosphorylated (pERK1/2) and therefore more active in NSCLC-NE than in SCLC (Figure 3C). DUSP6 protein was expressed in a subset of the ASCL1-high cells and some other NSCLC, but was not detected in the NEUROD1-high cells (H82, H524, HCC17)^63^ (Figure 3C). DUSP6 mRNA varies among the cell lines independently of the main transcription factor, as we observed in RNAseq data obtained from 247 lung cell lines (data not shown). However, DUSP6 protein expression levels are consistently high in the tested ASCL1-high lines, while it is not detected in the NeuroD1-high lines. DUSP6 protein decayed with a time course similar to that of ASCL1 (Figure S1B). Thus, we chose to evaluate the impact of manipulating DUSP6 expression on survival of these cells.

### DUSP6 inhibition enhances ERK signaling

The observation that ERK1/2 activities are relatively low in ASCL1-high cells suggested that molecules such as DUSP6 are induced to restrict ERK kinase activity by dephosphorylating and inactivating them. To test this possibility further, SCLC H889 cells were stimulated with serum, which usually is a strong ERK1/2 activator. Suprisingly, serum stimulation alone caused only a modest and transient increase in ERK1/2 activity measured as pERK (Figure 5A). On the other hand, serum stimulation of cells pretreated for 3 hours with the small molecule DUSP6 inhibitor BCI^64–67^ resulted in greater ERK1/2 activation that persisted for several hours, consistent with the idea that DUSP6 suppresses growth factor-induced activation of ERK1/2 in these cells (Figure 5A). A similar result was observed in NE-NSCLC HCC1833 (Figure 5B). DUSP6 is primarily localized to the cytoplasm and can act as a scaffold to retain ERK2 in this cellular compartment^68–70^. In examining HCC1833 cells treated with the DUSP6 inhibitor, we found that the localization of pERK changed from primarily cytoplasmic to strongly nuclear (Figure 5C, Figure S2A), indicating that inhibition of DUSP6 enabled pERK1/2 to concentrate in the nucleus. The difference in localization was also detected in H889 cells (Figure S2B).

We examined short term effects of inhibiting MEK1/2 with PD03 to prevent ERK1/2 activation and of inhibiting DUSP6 with BCI on signaling and expression of DUSP6 in ASCL1-high, H889 and HCC1833, and a NEUROD1-high cell line H82 (Figure S2C). Exposure to PD03 suppressed residual ERK1/2 activity, but increased MEK1/2 activity in the ASCL1-expressing lines, as deduced from elevated phosphorylation of the activation loop sites (pMEK1/2, pS217, pS221). Increased pMEK1/2 is indicative of reduced negative feedback from ERK1/2 to upstream components in the pathway^71^. The relative amount of the faster migrating DUSP6 band was increased by PD03 (Figure S2C), most likely indicating a decrease in DUSP6 phosphorylation; this would be expected if DUSP6 is no longer phosphorylated by ERK1/2^68, 72, 73^. The Schering-Plough ERK inhibitor (SCH) caused effects similar but smaller than effects of PD03 on DUSP6 and pMEK1/2^74^ (Figure S2C). Longer times of BCI exposure decreased the amounts of DUSP6 and ASCL1 that were detected and also caused activation of the related MAPK, c-Jun N-terminal kinase (JNK) (Figure S2C), suggesting that high concentrations of the inhibitor caused a stress response. Together these results suggest that, in SCLC and NSCLC-NE lung cancers, a small amount of residual ERK1/2 activity is necessary, but that this activity is insufficient to induce strong feedback inhibition of ASCL1 expression using upstream steps in the activation pathway. With pharmacologic perturbation (e.g. with a MEK or DUSP6 inhibitor) the role this residual activity is playing and the importance of DUSP6 in its regulation is exposed.

### Pharmacological inhibition of DUSP6 and genetic knockout affect proliferation and survival in ASCL1-high cells

The DUSP6 inhibitor BCI caused a decrease in attached cells and a loss of ASCL1 expression (Figure 4A). Decrease in ASCL1 protein levels were apparent in the time course of Figure 5A. The DUSP6 inhibitor also induced caspase cleavage within 30 min (Figure 4B). These results support the conclusion that DUSP6 inhibition, with BCI, leads to pERK activity, which in turn reduces ASCL1 expression which is associated with development of apoptosis. Consistent with apoptosis induction, exposure of HCC1833 cells to BCI for 2 weeks prevented formation of soft agar colonies (Figure S3A). However, BCI also reduced colony formation of NEUROD1-high H82 cells, consistent with some DUSP6-independent toxicity or off-target effects. Survival of ASCL1-expressing cells in the presence of BCI was reduced relative to the NEUROD1-high H82 line (Figure S3C).

**Figure 4.**
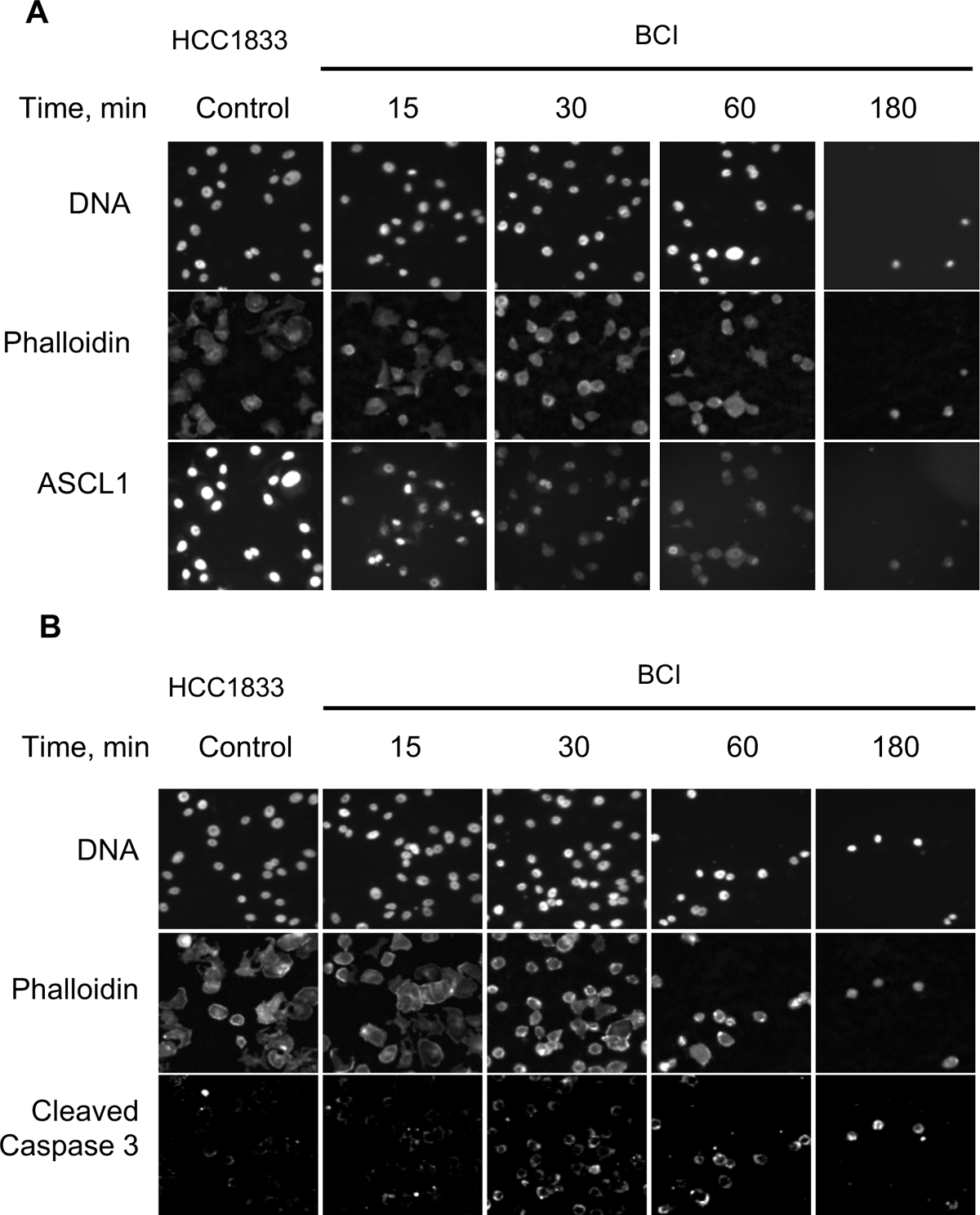
Inhibition of DUSP6 reduces ASCL1 and induces caspase activation. HCC1833 cells were exposed to 2.5 μM DUSP6 inhibitor BCI over a 3 hour time course. The cells were stained with DAPI to show DNA and phalloidin to show cell boundaries. Immunostaining shows ASCL1 (**A**) and cleaved caspase (**B**). Fields are shown from one of two replicates.

An obvious approach is to CRISPR edit or knockdown DUSP6 with functional genomics to determine the effects on SCLC survival and ASCL1 expression. However, we found using multiple methods to edit the *DUSP6* gene in H889 produced no live cells. In the case of ASCL1 high HCC1833 cells, DUSP6 gene knockout was successfully achieved, which led to a significant reduction in proliferation, mirrored by DUSP6 pharmacological inhibition by BCI in the non-modified cells (Figure 5D). Notably, addition of BCI to the DUSP6 KO cells did not further reduce growth (Figure 5D). Consistent with proliferation assays, viability of HCC1833 cells was reduced by DUSP6 knockout; the effect of the KO was greater than treatment with BCI, which did not further decrease viability compared to the KO alone (Figure 5E). Importantly, these experiments indicate the findings with BCI appear to be predominantly on-target effects of this drug at the low concentration (0.5 µM) used here. Of great importance, we discovered the tumor cells rapidly adapted to DUSP6 gene editing and recovered their ability to grow after a few passages (Figure S3E). However, this growth restoration was not due to a reexpression of DUSP6 as we observed by western blot analysis (Figure S3F). Further anlysis of this interesting observation will define the adaptation mechanism to the lack of DUSP6 in future work.

**Figure 5.**
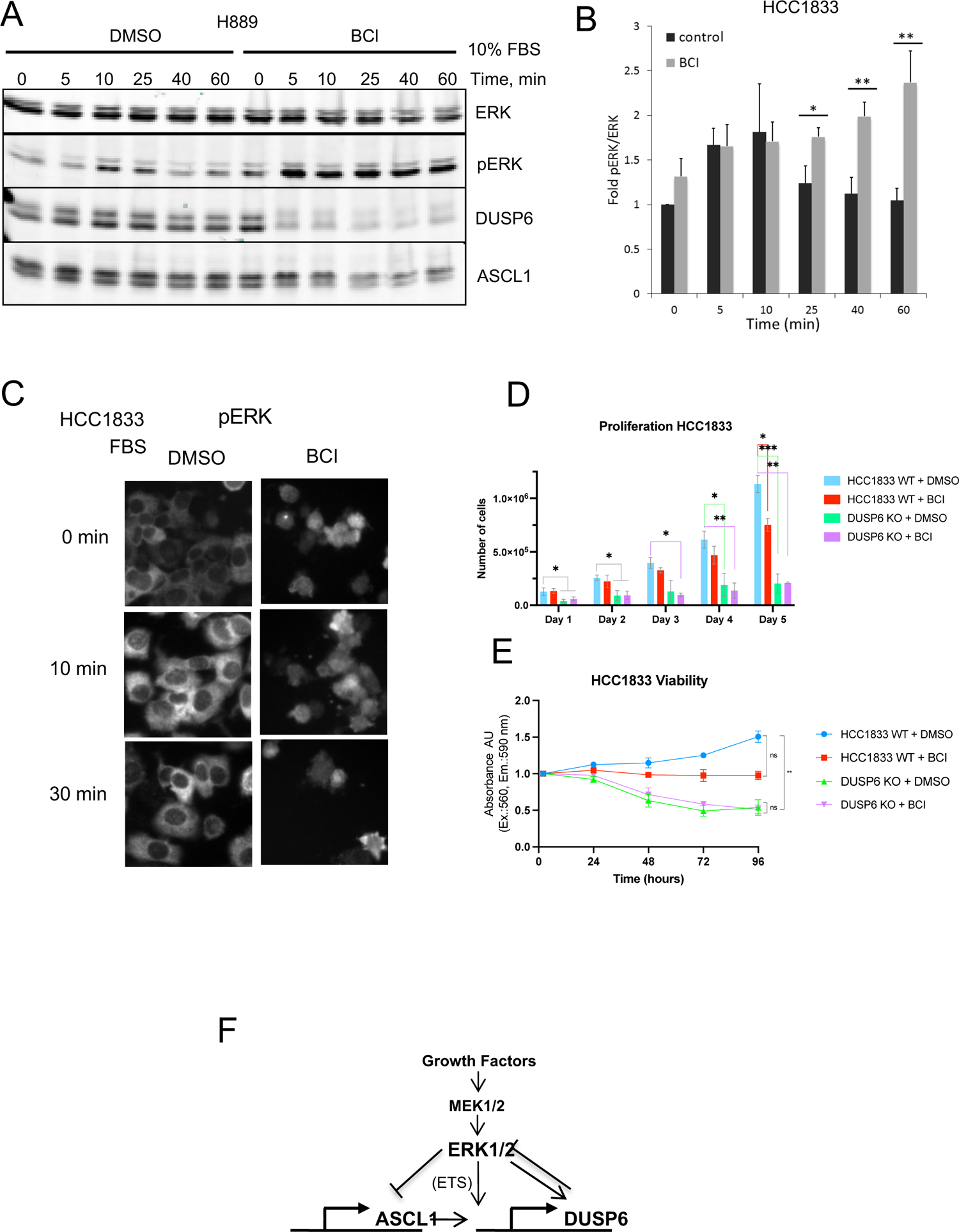
Inhibition of DUSP6 increases ERK activity and nuclear localization. **A.** H889 cells were deprived of serum and treated with 50 μM BCI or DMSO for 3 hours prior to the addition of 10% fetal bovine serum. ERK and pERK, ASCL1 and DUSP6 were measured by immunoblotting over the time course. N=2. **B.** HCC1833 were treated as in A. Immunoblots of ERK and pERK were quantified. N=3; * p<0.05, ** p<0.01. **C.** HCC1833 cells were pretreated with 2.5 μM BCI for 3 hours and stimulated with 20% FBS over the indicated times. The localization of pERK was assessed by immunofluorescence. Fields are shown from one of two replicates. **D.** Proliferation assay in HCC1833 WT and DUSP6 KO cells -/+ BCI. 0.5 µM BCI was applied in day 1 and refreshed daily over 5 days. Cells were trypsinized and two technical replicates were counted every 24 h. N=3 with 2 replicates. Statistical analysis: two-way ANOVA multiple comparisons. **E.** Viability assay in HCC1833 WT and DUSP6 KO cells -/+ BCI by CellTiter-Blue. Cells treated as in D for 4 days. N=3 with 3 replicates. Statistical analysis: one way ANOVA multiple comparisons. **F.** Model proposed to account for interactions between ASCL1 and ERK1/2.

From these results, we propose a model in which ERK interferes with expression of ASCL1 at the level of transcription and ASCL1 prevents a reduction in its own expression by inducing transcription of genes encoding proteins that interfere with ERK signaling, e.g., DUSP6 (Figure 5F).

## Discussion

In some tumor cell contexts, ASCL1 is associated with increased transcription of *DUSP6*, a phosphatase that selectively inactivates ERK1/2 MAPKs. ASCL1 knockdown resulted in enhanced ERK1/2 activity, confirming that ASCL1 can induce a program including *DUSP6* to suppress ERK1/2 function (see Figure 5F). DUSP6 is not the universal mechanism for suppressing ERK1/2 in ASCL1-high tumors, however, as it is not expressed in all cells with low ERK1/2 activity, but rather occurs in only a subset of these cancers^42^. Other mechanisms must function in different tumors. Suggested by our RNA-seq analysis, sprouty proteins such as SPRY4 that suppress activation of the ERK pathway by blocking signals downstream of receptor tyrosine kinases^75^, are also likely to be in play in other ERK1/2-sensitive tumor cells.

ASCL1 expression was decreased by extracellular ligands and a small molecule inhibitor of DUSP6 that stimulated ERK1/2 to varying extents in ASCL1-high SCLC and NSCLC-NE cell lines. The consequences included slowing of the cell cycle and increased cell death. Our results substantiate earlier studies showing that ERK1/2 activation by a Raf fusion protein interfered with proliferation and survival of SCLC lines^39^ and similar observations were made by ERK1/2 induction by the constitutively active mutant MEK DD, an active form of a direct activator of ERK^42^. Because ERK1/2 blockade with a MEK inhibitor enhanced expression of *ASCL1* mRNA, ERK1/2 apparently can suppress *ASCL1* transcription in these cancer cells. Besides suppression by the Notch signaling pathway and the transcriptional repressor REST, few mechanisms regulating *ASCL1* expression are understood^76, 77^. Many ETS factors including ELK1 and ETV5 are ERK1/2 substrates, and it seems possible that these factors may mediate ERK1/2-dependent suppression of ASCL1 transcription^78, 79^. As observed in this study by RNAseq, ETV5 expression was downregulated in H889 by exposure to the MEKi PD03. In line with this observation, it was recently described that ETV5 expression was specifically upregulated by ERK pathway activation via CBP/p300, consistent with downregulation of neuroendocrine transcription factors in some SCLC lines^41^. Determining upstream regulators of *ASCL1* may uncover new routes to therapy for pulmonary neuroendocrine tumors.

PMA was viewed as a tumor-promoting agent for many years due to activation of PKC and downstream pathways such as ERK1/2^80^. Context-dependent tumor suppressive effects in cell lines and cancer models have been attributed to PMA^81, 82^. More recently PKCs have been shown unequivocally to be tumor suppressors^83^. Because a dominant action of the ERK1/2 pathway is not stimulatory but instead inhibitory to SCLC and NSCLC-NE, it remains to be determined if PKCs may also have growth stimulatory effects in these neuroendocrine cancers, in contrast to their now-known tumor suppressive actions in many other cancer types.

BCI was identified in a zebrafish screen for DUSP6 inhibitors and is one of few small molecules with DUSP selectivity. As a result, BCI has been used in numerous studies to implicate the importance of DUSP6 in various types of cancer^64–67, 84–88^. In these studies and ours, one validation of BCI as a DUSP inhibitor was its ability to increase cellular pERK1/2. To provide evidence that additional actions of BCI can realistically be attributed to DUSP6, we compared the effects of BCI on a cell line that does not detectably express DUSP6 (H82). H82 cells not expressing DUSP6 were affected by BCI, but displayed reduced sensitivity to the compound compared to the DUSP6-expressing cells, confirming some on-target action of the molecule. Furthermore, BCI was tested in HCC1833 cells that were genetically edited to delete the target, DUSP6. This compound did not additionally affect proliferation or viability in cells already devoid of DUSP6. However, a reduction in both biological processes was observed upon BCI treatment of HCC1833 cells with intact DUSP6. In this study, BCI was used at different concentrations based on treatment duration and cell confluence. Nevertheless, because there is a lack of DUSP6 inhibitors with distinct chemical structures, where possible independent validation by genetic means should continue to be employed.

DUSP6, which is selective for ERK1/2 inactivation, can be viewed as a classic negative feedback regulator of ERK1/2 signaling in that activation of these kinases can promote transcription of *DUSP6* through ETS family transcription factors that are ERK substrates^56, 57^. However, interactions of ERK1/2 and DUSP6 are not that simple. ERK1/2 have been reported to influence stability of *DUSP6* mRNA, and can phosphorylate multiple sites on DUSP6, some inhibiting and some promoting its degradation^56, 73, 89^. Furthermore, DUSP6 requires allosteric activation by binding to a noncatalytic docking site on ERK1/2, independent of ERK kinase activity^90^. Thus, ASCL1, DUSP6, and ERK1/2 form a multicomponent complex feedback pathway in neuroendocrine lung cancers. The answer to an important question is, however, still incomplete and that is how does ASCL1 skew the balance point so strongly toward suppression of ERK1/2 activity?

As a MAPK phosphatase, the most important function of DUSP6 is assumed to be dephosphorylating and inactivating ERK1/2. In HCC1833 cells, inhibition of DUSP6 resulted in the translocation of ERK1/2 to the nucleus, raising the possibility that anchoring ERK1/2 in the cytoplasm is also an important survival-promoting action of DUSP6. The cytoplasmic retention of ERK1/2 by DUSP6 may prevent nuclear actions that interfere with survival of SCLC and NSCLC-NE. In addition to suppressing *ASCL1* transcription, likely mediated by phosphorylation of transcriptional repressors, nuclear concentration of ERK1/2 may facilitate possible direct effects of ERK1/2 on transcriptional repression^91, 92^ or other events that impair survival of neuroendocrine tumor types. DUSP6 and ETV5 were found essential for oncogenic transformation of pre B-cell acute lymphoblastic leukemia^67^. Interactions among these factors has been delved into more deeply in an excellent recent study in lung cancers^41^.

Although DUSP6 is now viewed as a tumor promoter in multiple neuroendocrine tumors^93, 94^, in several other cancer types including some forms of NSCLC, strong evidence indicates that DUSP6 is a tumor suppressor^69, 88, 93, 95–97^. DUSP6 has been identified as a predictive marker for sensitivity of tumors to MEK1/2 inhibitors^98^ and is selectively downregulated by a long noncoding RNA that activates ERK1/2 in cutaneous squamous cell carcinoma^99^, for example. DUSP6 has also been identified as a suppressor of toxicity induced by a combination of EGF receptor and KRAS mutations, thereby facilitating tumor growth^88^.

We conclude that controlling ERK1/2 activity, either by neutralizing ligand activation or by maintaining activity within a limited range, enhances survival of certain neuroendocrine cancers that benefit from expression of ASCL1, recently also reported in other studies^41, 42^. As in NSCLC-NE like HCC1833, inhibitory effects of high ERK activity have also been found in melanoma and B-cell leukemia, for example^67, 100, 101^. The apparently paradoxical observations discussed above highlight the fact that ERK1/2 have many potential functions, based on their extensive and diverse substrates and array of control mechanisms, that can be tuned or adapted to provide contextually appropriate cell functions^102^. That same adaptability makes them uniquely exploitable in disease. While ERK1/2 apparently contribute positively to growth and survival of SCLC and other neuroendocrine cancers^43–47^, these kinases also have the potential to limit the proliferation and survival of these cancers if their activities escape rather narrow disease-specified limits.

### Limitations of the study

Genetic disruption of DUSP6 in H889 failed despite multiple efforts. This implies the importance of this protein for survival of these cells. In contrast, knockout was feasible in HCC1833 which showed a slower proliferation rate compared to the unmodified cell line along with a slight decrease in viability. Nevertheless, cells adapted to DUSP6 loss after a few passages, apparently by the upregulation of alternative genes or hyperactivation of secondary pathways compensating for the silenced gene which results in survival. DUSP6 may serve as a potential therapeutic target only in those ASCL1 lung tumors that express it. However, the possibility exists that other negative regulators would be involved in maintaining ERK activity within low optimal levels in other tumors.

## Author contribution

Conceptualization, MHC, JEJ, JDM; Methodology, MHC, JEJ; Investigation, AMV, SE, AA, CW, KM, MP; Formal Analysis, CW, RKK, LG; Writing – Original Draft, MHC; Writing –Review & Editing, AMV, JDM, JEJ; Funding Acquisition, MHC, JEJ, JDM; Resources, JDM, AG, JEJ, MHC; Supervision, MHC, JEJ.

## Acknowledgments

The authors thank Ankita Jaykumar, Ji-Ung Jung, Mike Kalwat (Indiana Biosciences Research Institute) and other current and former members of the Cobb, Johnson, and Minna labs for valuable suggestions, Gray Pearson (Georgetown University) for comments on initial figures, and Dionne Ware for administrative assistance. These studies were supported by CPRIT RP110383 and RP140143 to MHC and JEJ, 1U01CA213338 to JDM, Welch Foundation grant I1243 to MHC, the Hamon Center NCI Spore Grant in Lung Cancer P50CA70907 to JDM, Mary Kay Fellowship to AM-V, and the Simmons Comprehensive Cancer Center grant (P30 CA142543) for assistance from the UTSW microscopy core.

---

**Table S1.** RNA-seq of H889 cells treated with PD0325901 at 100 nM and 1 μM.

**Table S2.** Cell lines used in this study.

## STAR Methods

### RESOURCE AVAILABILITY

### Lead contact

Further information and requests for resources and reagents should be directed to and will be fulfilled by the lead contact, Melanie Cobb.

### Materials availability

This study did not generate new unique reagents.

### Data and code availability

- RNA-seq data is available in this paper’s supplemental information. Western blot and microscopy data reported in this paper will be shared by the lead contact upon request.
- This paper does not report original code.
- Any additional information required to reanalyze the data reported in this paper is available from the lead contact upon request.

## EXPERIMENTAL MODEL AND STUDY PARTICIPANT DETAILS

### Cell culture and cell lines

Cell lines used in this study, sex is indicated in brackets (F: female, M: male): H889 (F), HCC1833 (F), H1184 (M), H128 (M), H2107 (M), H2029 (F), H82 (M), H524 (M), H1755 (F), HCC4018 (F), HCC2374 (F), H460 (M), H1993 (F), H2882 (F), H2087 (M), H774 (M) and H720 (M). All lung cancer cell lines used in this study were provided by the Hamon Center for Therapeutic Oncology (University of Texas Southwestern Medical Center) and were fingerprinted (PowerPlex 1.2 Kit, Promega) and mycoplasma-free (e-Myco Kit, Boca Scientific). Cells were maintained in RPMI-1640 (Life Technologies Inc.) supplemented with 5% fetal bovine serum (FBS) without antibiotics at 37°C in a humidified atmosphere containing 5% CO_2_.

### Treatments

Cycloheximide (Sigma-Aldrich); DUSP6 inhibitor, BCI hydrochloride (Sigma-Aldrich); phorbol myristate acetate (Sigma-Aldrich); chloroquine diphosphate salt (Sigma-Aldrich); ERK inhibitor, SCH772984 (MedChemExpress); MEK inhibitors, PD0325901 (MedChemExpress) and AZD6244 (Selleck Chem); PKC activator, phorbol myristate acetate (PMA) (Sigma-Aldrich). Drugs were added to the medium as described in the text and figure legends.

## METHOD DETAILS

### Immunoblotting

Cells were rinsed with ice-cold phosphate-buffered saline (PBS) and lysed in 50 mM Hepes pH 7.5, 150 mM NaCl, 1.5 mM MgCl_2_, 1 mM EDTA, 10% glycerol, 0.2% NaVO4, 100 mM NaF, 50 mM β-glycerol phosphate, 0.2% Triton ×100, and protease inhibitors all from Sigma: pepstatinA (0.4 µg/ml) (P4265), tosyl lysine chloromethyl ketone (4 µg/ml) (T7254), tosyl arginine methyl ester (4 µg/ml) (T4626), benzoyl arginine methyl ester (4 µg/ml) (B4256), soybean trypsin inhibitor (4 µg/ml) (T6522), and leupeptin (0.4 µg/ml) (L2023). Unless indicated, lysates were clarified by sedimentation for 10 min at 13,000 g at 4°C. Laemmli electrophoresis loading buffer (5X) was added to supernatants and the material was heated at 90°C for 3 min. Proteins were separated by polyacrylamide gel electrophoresis and transferred to nitrocellulose for immunoblotting. Imaging of immunoblots was performed using the LiCor Odyssey system. For experiments in Figures 1A and 5, HCC1833 cells were washed twice in cold PBS, trypsinized, and snap frozen. Lysates were prepared in M-PER Mammalian Protein Extraction Reagent (Thermo Scientific, Cat#78501) with β-mercaptoethanol (0.5 mM) and boiled. Immunoblots used horseradish peroxidase-conjugated secondary antibody detected by enhanced chemiluminescence (Thermo Scientific).

### Immunofluorescence microscopy

Cells were plated on Falcon 96-well culture plates, (Corning Cat#353219) coated with growth factor-reduced matrigel (GFR) (Fisher Scientific Cat#CB-40230) 24 h before treatment. After treatments indicated in the text, cells were washed twice with PBS and fixed with 4% paraformaldehyde (vol/vol) in PBS. Cells were permeabilized with 0.2% Triton X-100 in PBS for 15 min at RT, incubated with 100 mM glycine in PBS for 20 min at RT, then washed 3 times for 10 min with PBS, incubated with 10% normal goat serum (vol/vol) at room temperature for 30 min, and then with primary antibodies at 4°C overnight. Cells were washed 3 times with PBS containing 0.1% Tween, and incubated with Alexafluor-conjugated secondary antibodies, Alexa Fluor 546 goat anti-mouse (A11030) and Alexa Fluor 488 goat anti-rabbit (A11034), at room temperature for 2 h, washed as above, mounted with DAPI Fluoromount-G (Southern Biotech #0100-20) and imaged on a BD Pathway Bioimager 855 microscope.

### Quantitative RT-PCR

RNA was extracted using TRI-Reagent (NC9330796, Fisher Scientific). cDNA was generated with an iScript cDNA synthesis kit (BioRad, Cat#1708890). qPCR was done using the iTaq Universal SYBR Green Supermix (Biorad, Cat#172-5125). Primers for ASCL1: Forward Primer: CGCGGCCAACAAGAAGATG; Reverse Primer: CGACGAGTAGGATGAGACCG; for beta-actin: Forward: AGGTCATCACTATTGGCAACGA; Reverse: CACTTCATGATGGAATTGAATGTAGTT. Actin was employed as an internal reference to normalize input cDNA. PCR reactions were performed using the CFX96 Real-Time PCR Detection System and analyzed with Bio Rad CFX Manager3.1 software. For Figure 5, PCR used TaqMan Assay probes (Applied Biosystems) and RNA was isolated using the miRNeasy kit (Qiagen). iTaq Supermix with Rox (Bio-Rad), a premade formula containing iTaq DNA polymerase, optimized buffers, nucleotides and Rox passive detection dye, was used to perform the qPCR reaction. Reactions were run in triplicate wells on a 96-well plate in a 7300 Real Time PCR System (Applied Biosystems). 7300 System Software (Applied Biosystems) was used to derive Ct values and ΔCt values were calculated using GAPDH or 18s amplification as a control. DUSP6 and GAPDH TaqMan assays were utilized in this study (Applied Biosystems; Hs04329643_s1 and Hs02758991_g1).

### Cell Cycle Analysis

Phorbol 12-myristate 13-acetate (PMA) was purchased from Sigma-Aldrich. HBEC-3KT and HCC1833 cells were plated in 6-well plates at a density of 2×10^5^ cells per well and allowed to adhere for 24 h prior to treatment. PMA was added at a concentration of 1, 10, and 100 nM for 24 h after which cells were harvested for cell cycle analysis. DNA content was measured from lung cancer cells following treatment with PMA or DMSO to determine the cell cycle profile and measure apoptosis. Cells were fixed in 70% EtOH, added slowly during fixation to prevent clumping, for 15 min on ice or overnight at −20°C. Cells were incubated in buffered staining solution containing 0.05% Triton X-100, 0.1 mg/mL RNase A, and 50 µg/mL propidium iodide (PI) in PBS for 40 min at 37°C. Following incubation, 3 mL fresh PBS was added to quench the reaction, cells were spun at 1000 RPM for 5 min at room temperature, and then resuspendend in 0.5 mL PBS and analyzed using a FACScan or FACSCalibur flow cytometer. The Watson algorithm was performed in FlowJo software (Treestar) to determine the distribution and gating of cells in different states of DNA replication.

### RNA-seq

3 ml of H889 culture (7.5×10^5^ cells) were exposed to either with PD0325901 100nM (or 1µM) or DMSO (0.1%) for 18 h. Cells were pelleted and washed with PBS. Cell pellets were used for RNA purification using Invitrogen™PureLink™ RNA Mini Kit (Cat#12183018A). RNA sample (5- 10 µg RNA at 75-190 ng/µl concentration) were submitted for Library generation and RNA sequencing analysis to The McDermott Center Next Generation Sequencing (NGS) Core, UTSW, Dallas, Texas.

### ChIP-seq

Browser tracks were generated in the UCSC Browser with data from^17, 103^ (14,92) for the following cell lines: H1755, HCC4018 (NE-NSCLC), H128, H1184, H2107 (SCLC), and control cell lines H524 and H82 (ASCL1(-) SCLC).

### Liquid Colony Formation Assays

For anchorage-dependent colony formation assay cells were plated in 6-well plates. (E/Z)-BCI hydrochloride (Sigma-B4313) was added to both the medium and the matrix. Two weeks later, colonies were stained with 0.5% crystal violet.

### Cell proliferation assay

6×10^4^ WT or DUSP6 KO HCC1833 cells were seeded per well in a 24- well plate, one for each time point (1-5 days). BCI or DMSO were added every 24 h starting at day 1. At the chosen time, cells were trypsinized and, once detached, an equal volume of the cell suspension was added to the chamber of the counting slide (3 technical replicates) and counted with the TC20 automated cell counter (BioRad).

### Cell viability assay

CellTiter-Blue Reagent (Promega) was used to assess cell viability. 10^4^ WT or DUSP6 KO HCC1833 cells were seeded per well in 96-well plates (one per each time point: 24, 48, 72 and 96 h). 24 h post seeding, the reagent was added, and the absorbance (560_ex_/590_em_) was measured after a 1 h 30 min incubation. BCI or DMSO were added for the first time at 24 h, as indicated in Figure S3E, and the treatments and media were refreshed daily. Absorbance was measured over the 4 days with 3 technical replicates per condition. Results were analyzed according to manufacturer’s instructions.

### Transient siRNA Transfections

Lung cancer cell lines were optimized for transfection conditions in 6-well plates by monitoring lipid content and cell number, and measuring the proliferative differences between scrambled oligo control (Qiagen) and toxic control (Qiagen). For 6-well experiments, 3-5 μL RNAiMAX (Invitrogen) were added to 500 uL serum-free RPMI-1640 and incubated at room temperature for 5 min. 20 nM siRNA was mixed, plated dropwise in 6-well plates, and complexed for 20 min. 2×10^5^ cells supplemented with 5% were added on top of the mixture, and incubated at 37°C for 72 h prior to analysis. siRNAs were purchased from Qiagen, including siASCL1-1,-2,-3 (SI00062573, SI00062580, SI00062587), siLUC (SI03650353), and siSCR (SI03650325).

### CRISPR/Cas9 cell line generation

The short guide sequences (gDusp6-3C-Forward: caccGACGACTCGTATAGCTCCTG; gDusp6-3C-Reverse: aaacCAGGAGCTATACGAGTCGTC) were cloned into pSpCas9(BB)-2A-GFP (PX458) (Addgene) following the Sequence Cloning Protocol from Zhang Lab (https://www.addgene.org/crispr/zhang/). The constructs were transfected into HCC1833 cells using FuGeneHD reagent. Following transfection, cells were allowed to grow for 48 hours before sorting for GFP-positive cells. GFP-positive single cells were placed into an individual well of a 96-well plate. Cell colonies were formed in over 2 weeks. Individual colonies were then expanded and subsequently validated by testing for DUSP6 protein expression through Western blot analyses.

### Dose-response assay

H889 cells were tested for sensitivity to AZD6244. Cells were plated in 96 well plates at 10^4^ cells per well in 50 µl RPMI-1640 supplemented with 5% serum on day one. On day two, 50 µl of 2X drug was added in 8 doses using a four-fold dilution series in the same medium. For AZD6244 the top dose was 256 µM. On day five, 20 µl of CellTiter 96® AQueous One Solution Promega (Cat#G3580) was added to each well. The plates were incubated at 37°C for 1 h and then read on an absorbance reader at 490 nm. Control wells (cells with no drug) were normalized to 100% and each drug dose was expressed as a fraction of the control. 8 technical replicates per condition, N=2.

## QUANTIFICATION AND STATISTICAL ANALYSIS

Statistical analyses were performed using GraphPad Prism software version 9.5.1 (San Diego, CA). Results were expressed as means ± S.E.M determined from at least three independent experiments. Statistical significance was calculated using the t-test. Where indicated, statistical significance was assessed using one-way ANOVA or two-way ANOVA multiple comparisons. *P* values: \*\*\**p*<0.005, ** *p*<0.01, * *p*<0.05, ns (no significant).

## KEY RESOURCES TABLE

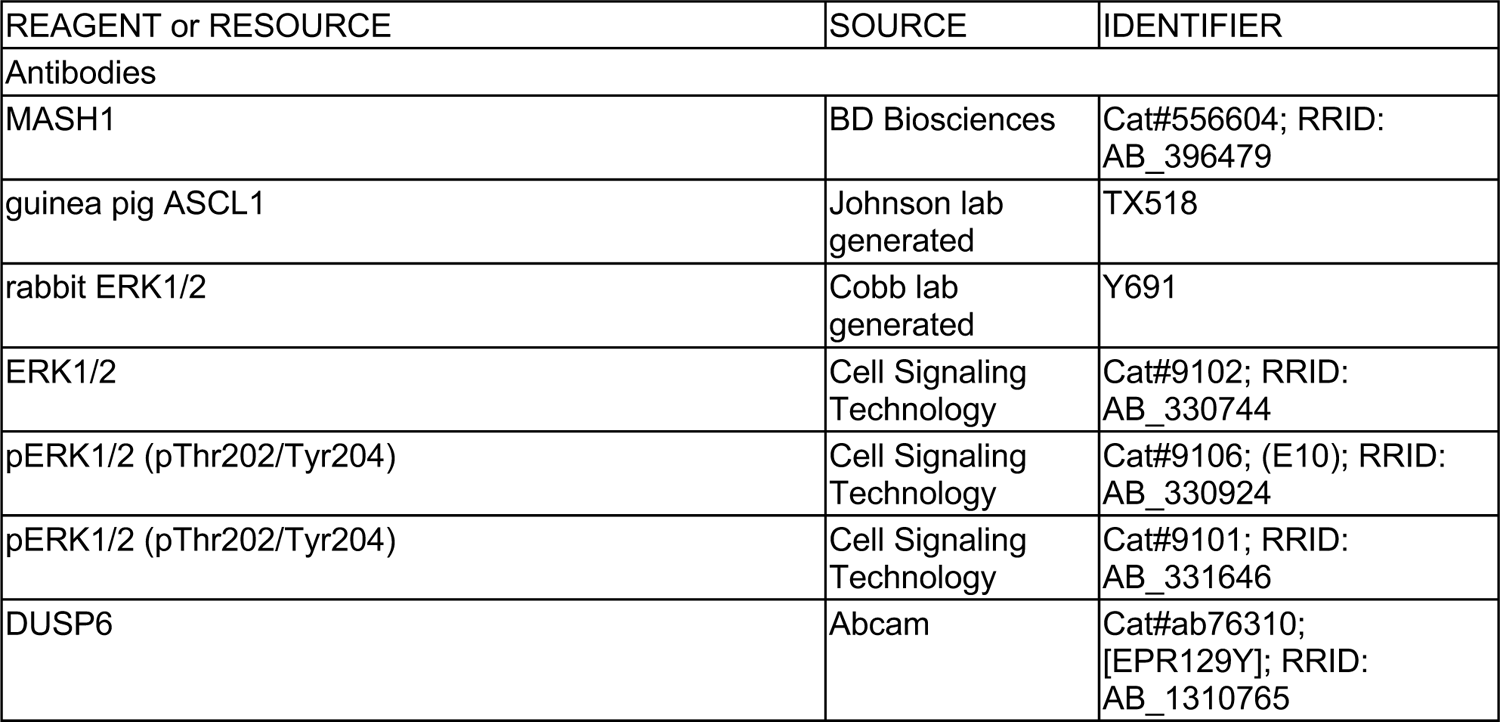
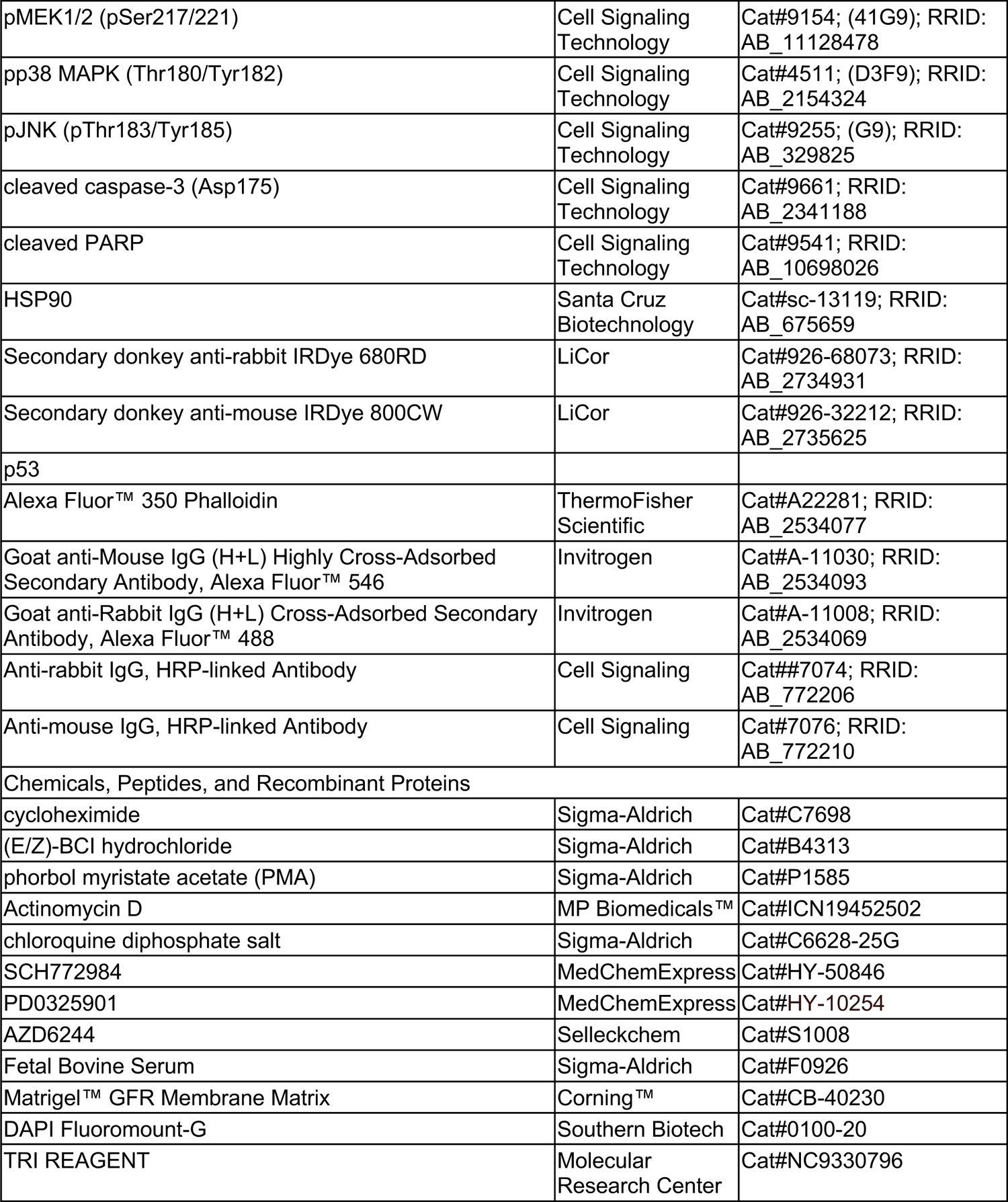
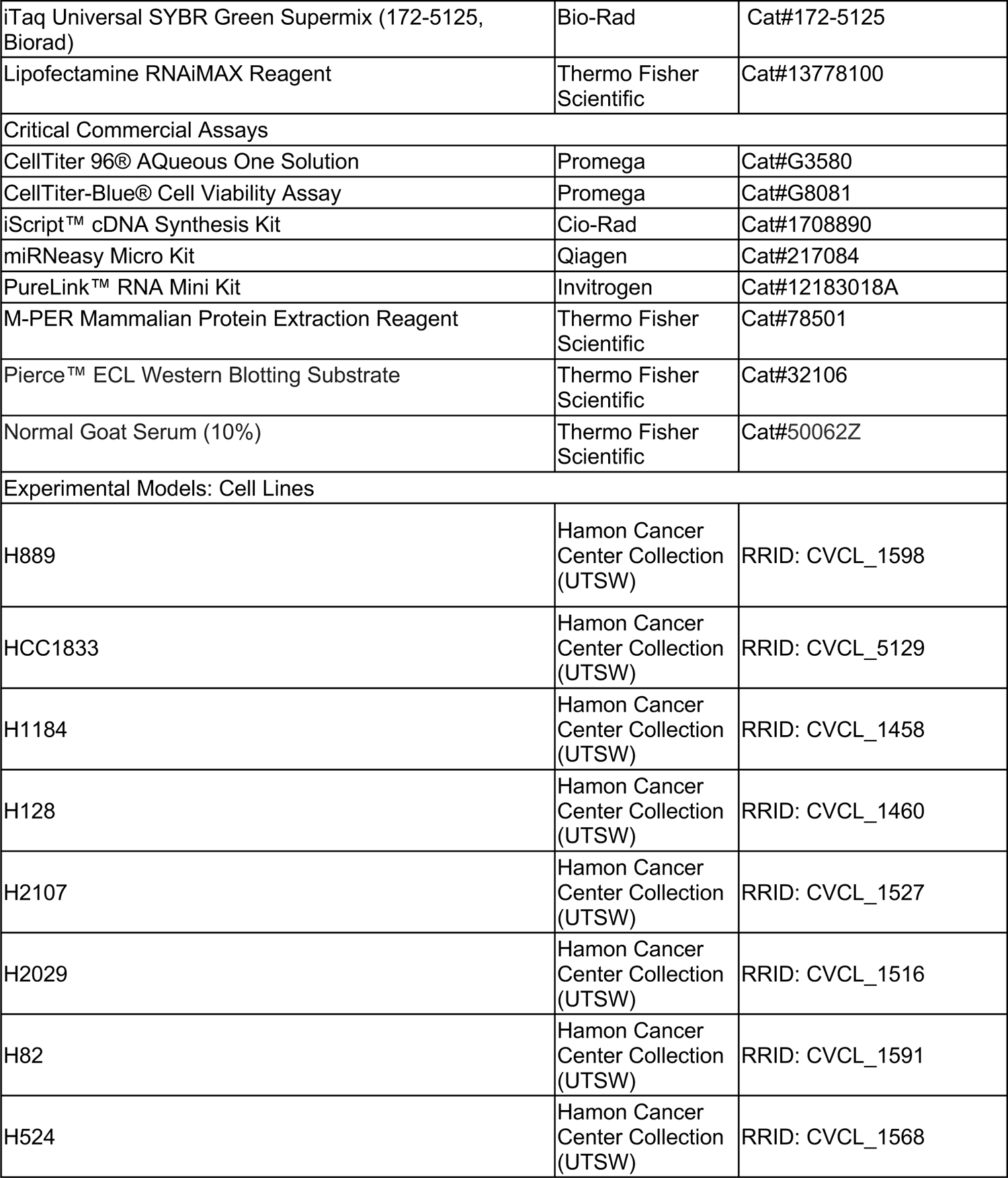
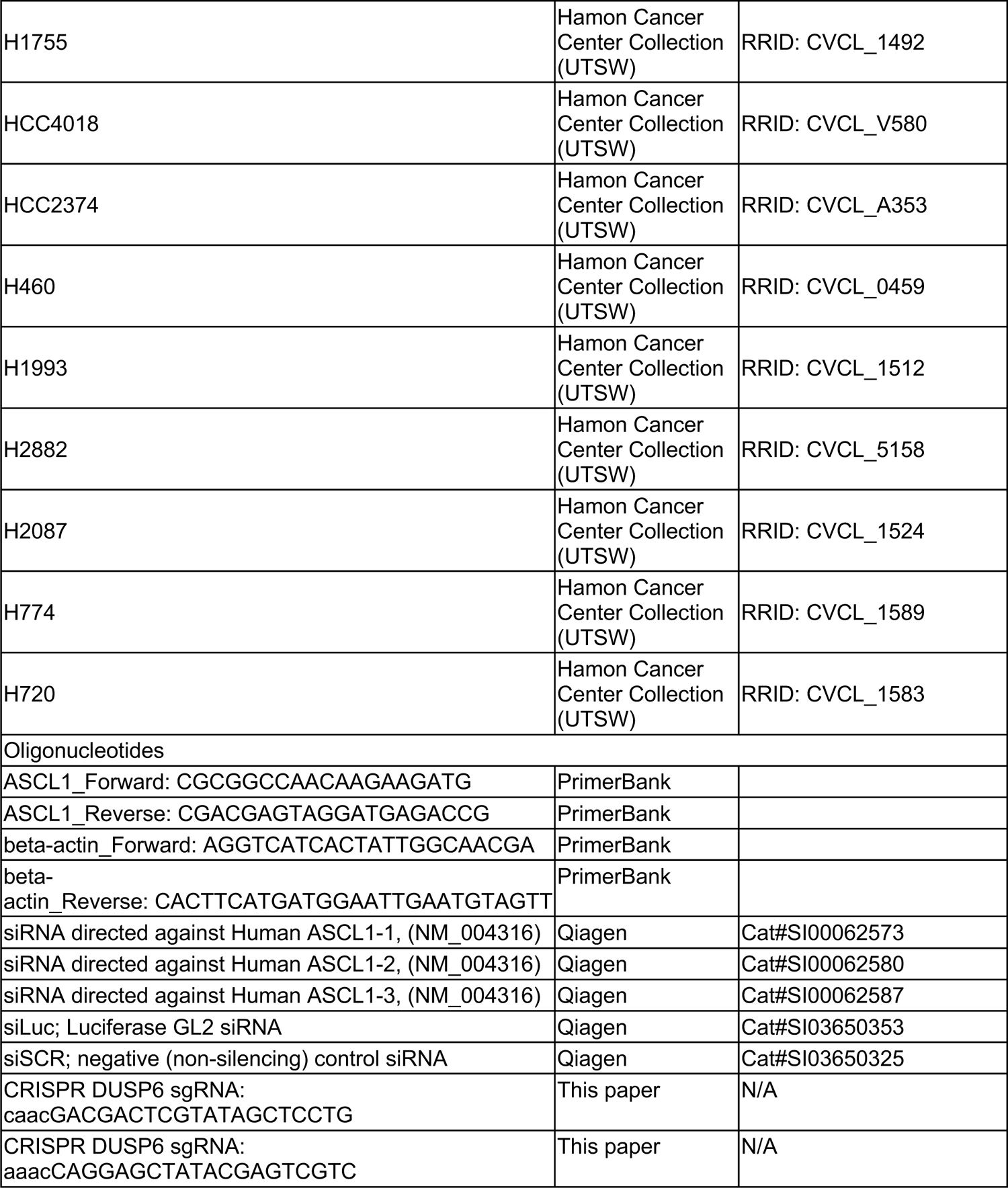
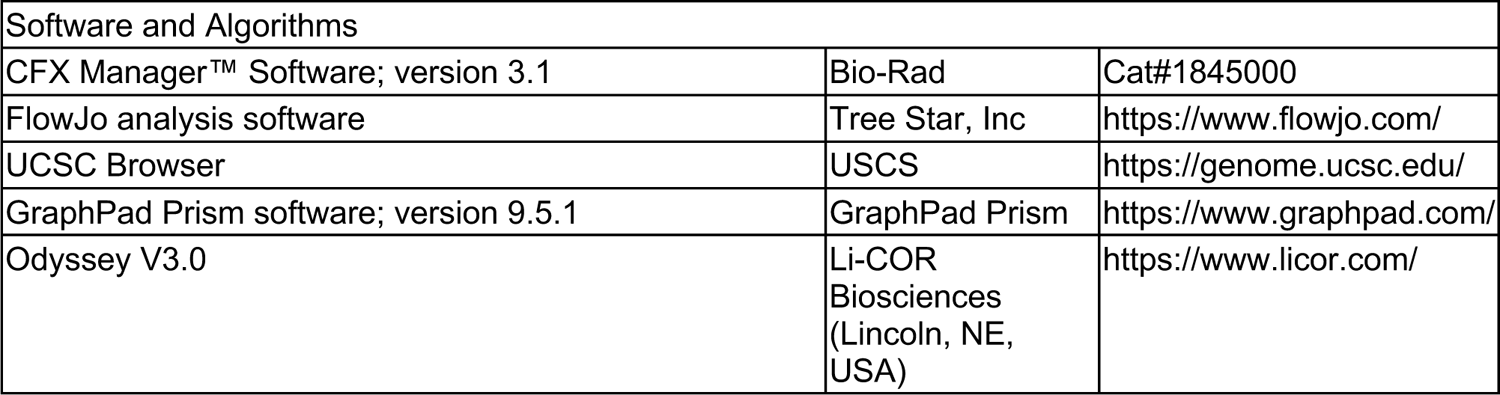

