## Supplemental Figures 1, 2 and 3 for "ASCL1-ERK1/2 Axis: ASCL1 restrains ERK1/2 via the dual specificity phosphatase DUSP6 to promote survival of a subset of neuroendocrine lung cancers"

### Slide 1
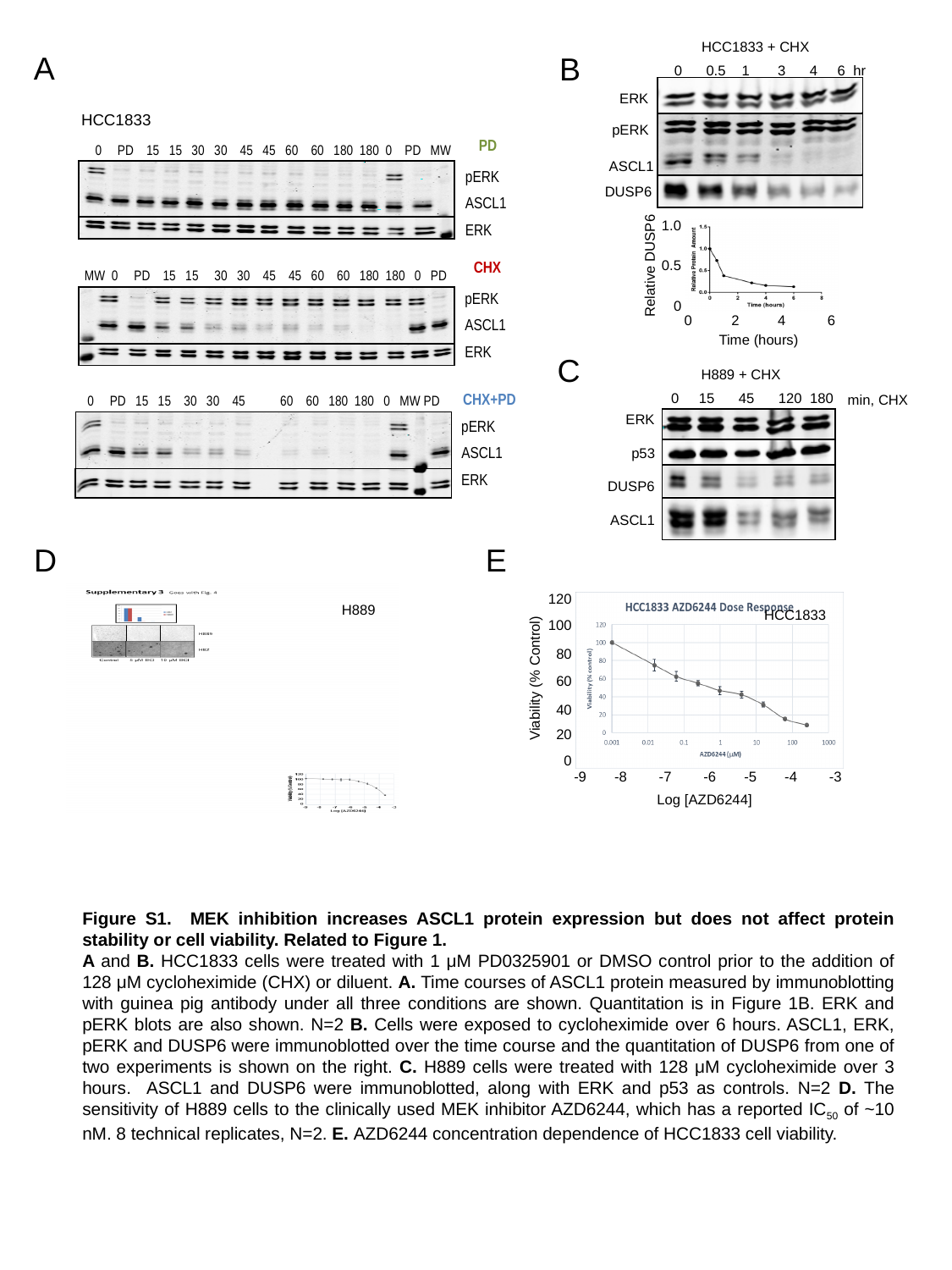

HCC1833 + CHX
 0 0.5 1 3 4 6 hr
ERK
pERK
ASCL1
DUSP6
A
B
HCC1833
PD
0 PD 15 15 30 30 45 45 60 60 180 180 0 PD MW
CHX
MW 0 PD 15 15 30 30 45 45 60 60 180 180 0 PD
CHX+PD
 0 PD 15 15 30 30 45 60 60 180 180 0 MW PD
pERK
ASCL1
ERK
pERK
ASCL1
ERK
pERK
ASCL1
ERK
1.0
0.5
Relative DUSP6
0
0
2
4
6
Time (hours)
C
H889 + CHX
 0 15 45 120 180
ERK
p53
DUSP6
ASCL1
min, CHX
D
E
100
0
120
20
80
Viability (% Control)
60
40
-9 -8 -7 -6 -5 -4 -3
Log [AZD6244]
HCC1833
H889
Figure S1. MEK inhibition increases ASCL1 protein expression but does not affect protein stability or cell viability. Related to Figure 1.
A and B. HCC1833 cells were treated with 1 μM PD0325901 or DMSO control prior to the addition of 128 μM cycloheximide (CHX) or diluent. A. Time courses of ASCL1 protein measured by immunoblotting with guinea pig antibody under all three conditions are shown. Quantitation is in Figure 1B. ERK and pERK blots are also shown. N=2 B. Cells were exposed to cycloheximide over 6 hours. ASCL1, ERK, pERK and DUSP6 were immunoblotted over the time course and the quantitation of DUSP6 from one of two experiments is shown on the right. C. H889 cells were treated with 128 μM cycloheximide over 3 hours. ASCL1 and DUSP6 were immunoblotted, along with ERK and p53 as controls. N=2 D. The sensitivity of H889 cells to the clinically used MEK inhibitor AZD6244, which has a reported IC_50_ of ~10 nM. 8 technical replicates, N=2. E. AZD6244 concentration dependence of HCC1833 cell viability.

### Slide 2
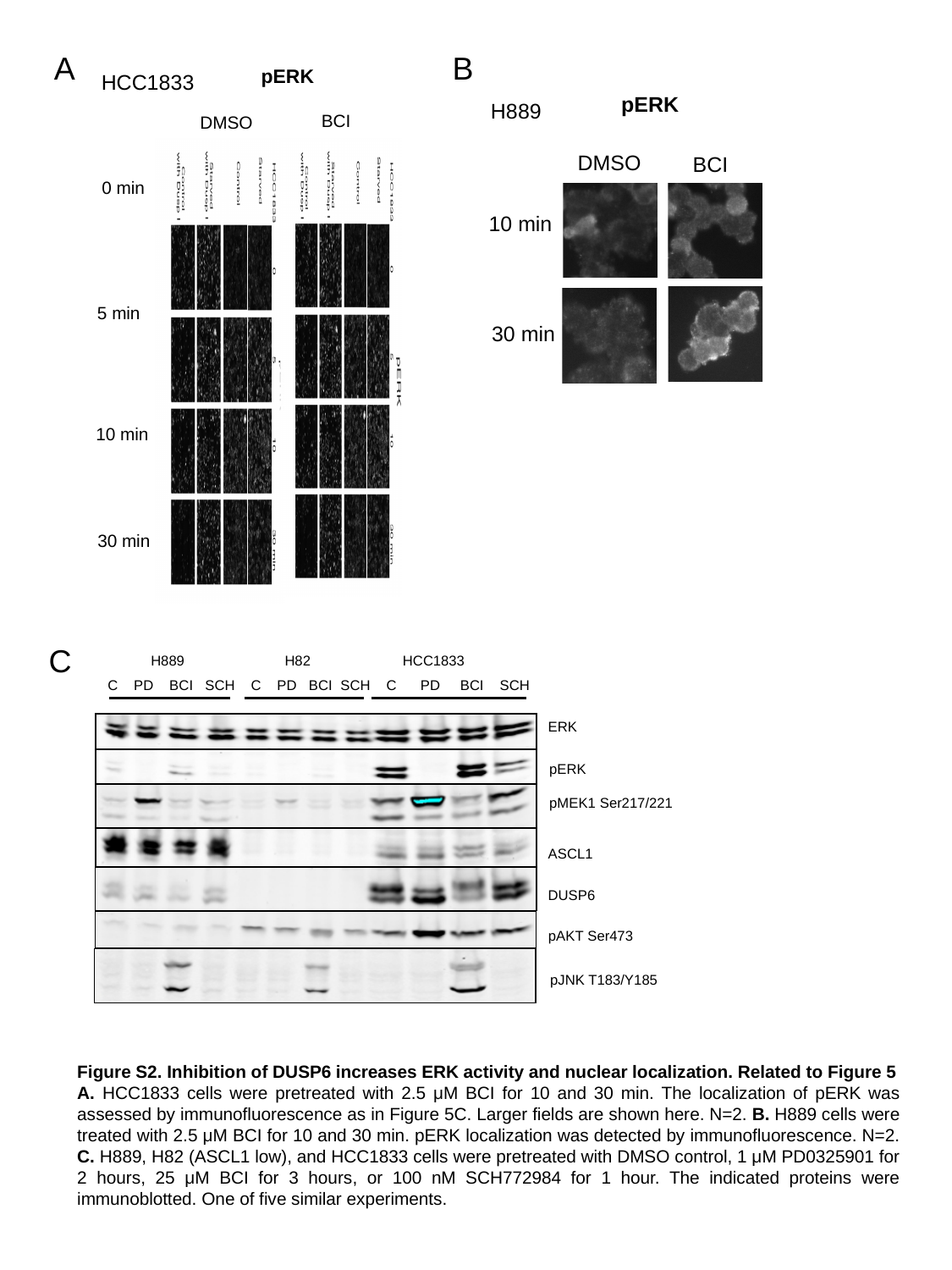

A
B
pERK
HCC1833
BCI
DMSO
0 min
5 min
10 min
30 min
pERK
H889
BCI
10 min
30 min
DMSO
C
H889 H82 HCC1833
C PD BCI SCH C PD BCI SCH C PD BCI SCH
ERK
pERK
pMEK1 Ser217/221
ASCL1
DUSP6
pAKT Ser473
pJNK T183/Y185
Figure S2. Inhibition of DUSP6 increases ERK activity and nuclear localization. Related to Figure 5
A. HCC1833 cells were pretreated with 2.5 μM BCI for 10 and 30 min. The localization of pERK was assessed by immunofluorescence as in Figure 5C. Larger fields are shown here. N=2. B. H889 cells were treated with 2.5 μM BCI for 10 and 30 min. pERK localization was detected by immunofluorescence. N=2. C. H889, H82 (ASCL1 low), and HCC1833 cells were pretreated with DMSO control, 1 μM PD0325901 for 2 hours, 25 μM BCI for 3 hours, or 100 nM SCH772984 for 1 hour. The indicated proteins were immunoblotted. One of five similar experiments.

### Slide 3
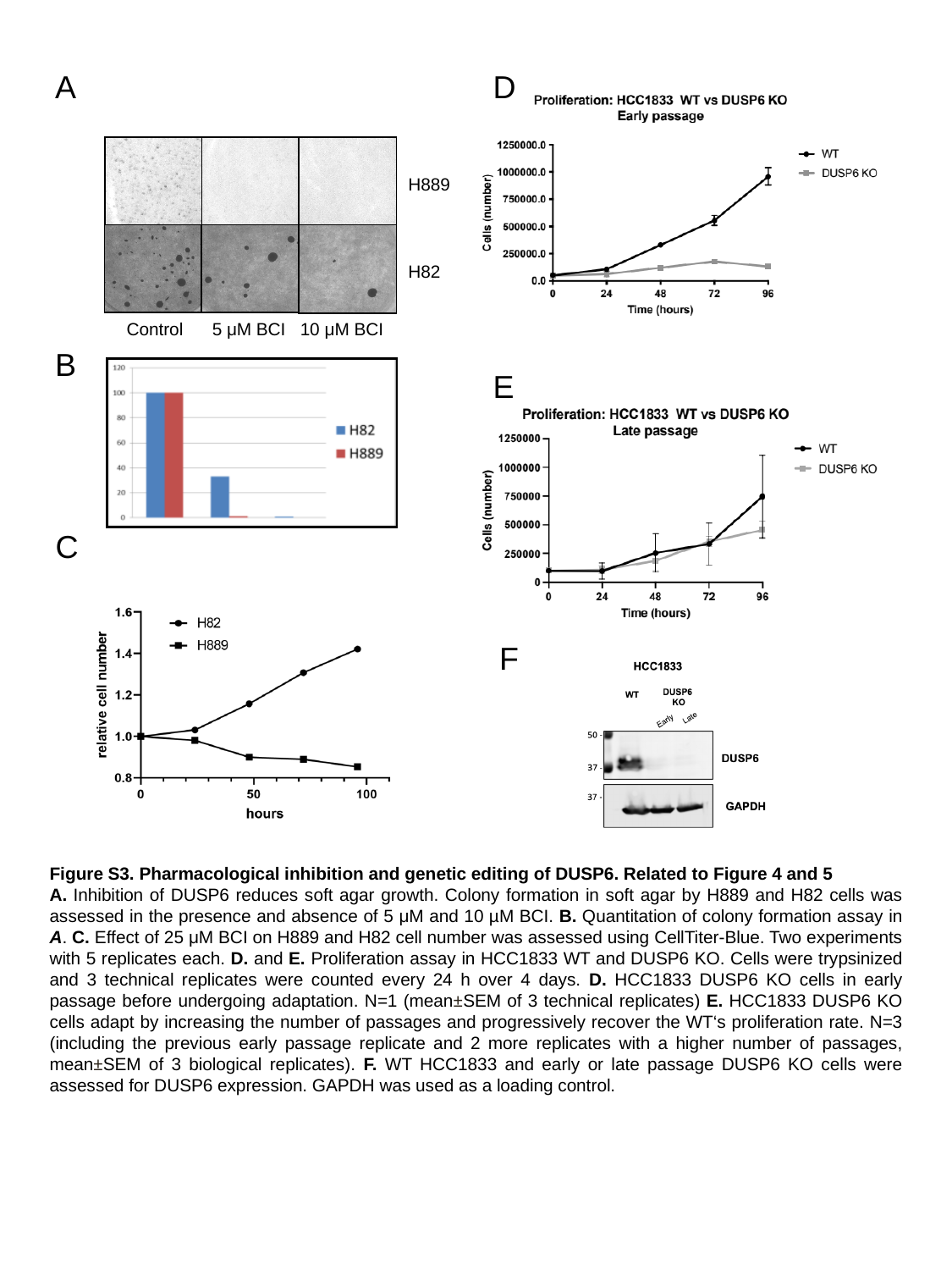

A
D
H889
H82
Control 5 μM BCI 10 μM BCI
B
E
C
F
Figure S3. Pharmacological inhibition and genetic editing of DUSP6. Related to Figure 4 and 5
A. Inhibition of DUSP6 reduces soft agar growth. Colony formation in soft agar by H889 and H82 cells was assessed in the presence and absence of 5 μM and 10 µM BCI. B. Quantitation of colony formation assay in A. C. Effect of 25 μM BCI on H889 and H82 cell number was assessed using CellTiter-Blue. Two experiments with 5 replicates each. D. and E. Proliferation assay in HCC1833 WT and DUSP6 KO. Cells were trypsinized and 3 technical replicates were counted every 24 h over 4 days. D. HCC1833 DUSP6 KO cells in early passage before undergoing adaptation. N=1 (mean±SEM of 3 technical replicates) E. HCC1833 DUSP6 KO cells adapt by increasing the number of passages and progressively recover the WT‘s proliferation rate. N=3 (including the previous early passage replicate and 2 more replicates with a higher number of passages, mean±SEM of 3 biological replicates). F. WT HCC1833 and early or late passage DUSP6 KO cells were assessed for DUSP6 expression. GAPDH was used as a loading control.
